# Embodied timing turns a robotic tug-of-war into cooperative transport

**DOI:** 10.64898/2026.10.26.762051

**Authors:** Atanu Chatterjee, Arthicha Srisuchinnawong, Poramate Manoonpong, Saad Bhamla

## Abstract

Living systems move loads with agents that pull against one another. Kinesin and dynein pull one vesicle, ants haul a dead insect, and antagonist muscles move a joint. Whether the timing of the forces can by itself determine which side moves the load is not well understood. Here, combining experiments, theory, and simulations of a tug-of-war between two matched teams of spring-coupled robots, we show that the relative timing of the pull forces breaks the symmetry and decides the contest. Teams pulling in unison or in strict alternation tie. Between these, the lagging team wins, most strongly near a quarter cycle, because the springs unload while the leading team coasts and the lagging team still pulls. In the model and in robot simulations, braking rather than coasting between pulls, once strong enough, shifts the advantage to the leading team. Winning and losing leave distinct signatures in each robot’s motor current. Using these local signals, we formulate an embodied decentralized phase-adaptation rule that enables transient leadership to switch between teams, allowing the collective to transport high-friction cargo and to navigate around obstacles towards a prescribed target without explicit communication.

## INTRODUCTION

Moving a shared cargo is not simply a matter of summing forces. A group must generate enough force to overcome resistance, but it must also coordinate those forces so that the load moves towards a useful target. Cooperative transport spans scales, from intracellular cargoes carried by teams of kinesin and dynein motors acting in opposite directions to ants hauling objects far larger than a single worker (Fig. 1A,B) [1–5]. In both cases, the load is not only the object being moved but also the medium through which the agents interact. A force applied by one agent changes the mechanical state experienced by the others, allowing the load itself to convey information about the collective state [6, 7].

**FIG. 1.**
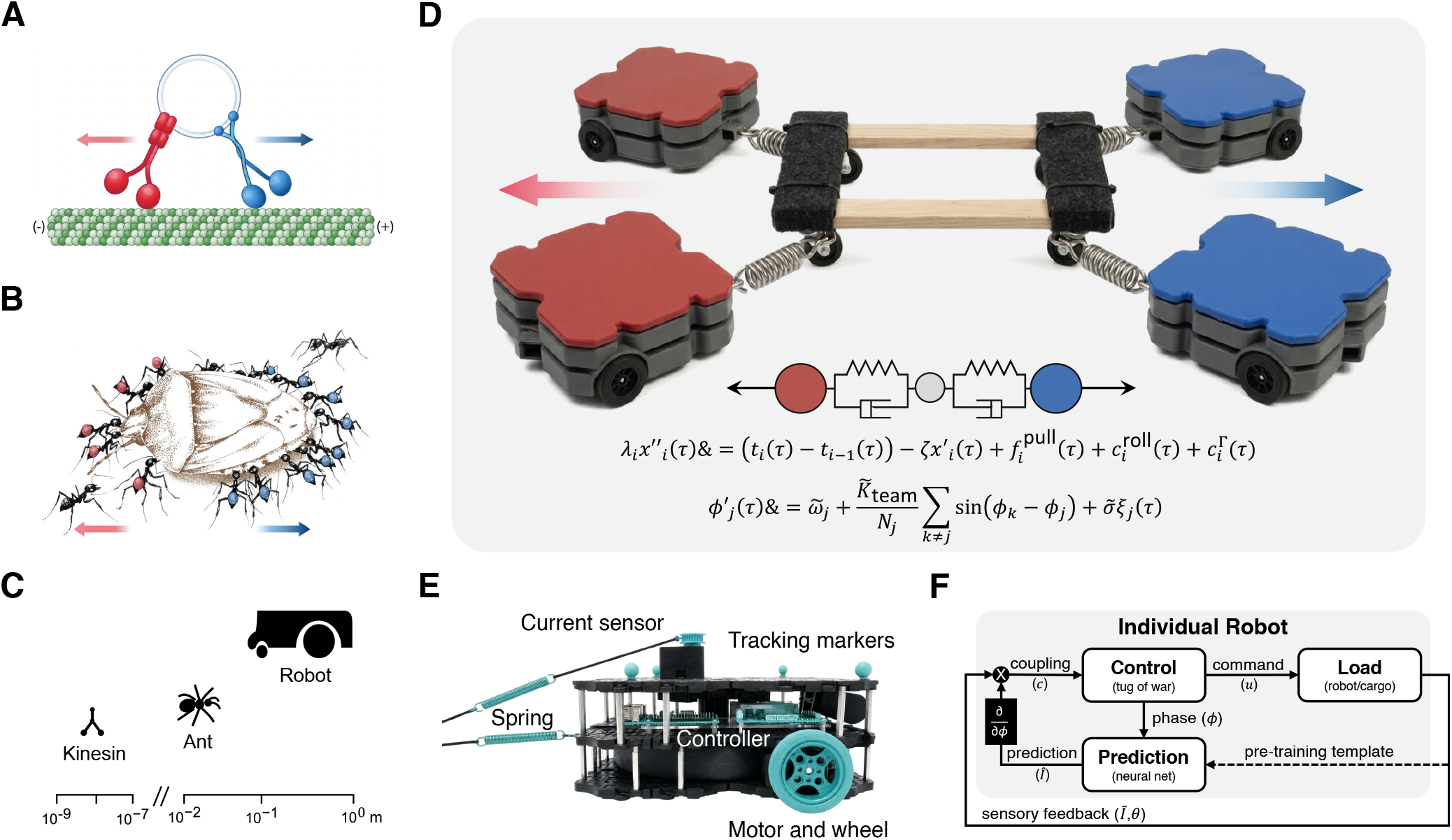
Competing forces in cooperative transport across scales. (A) A molecular scale tug-of-war emerges when kinesin and dynein motors transport a cargo along a microtubule. (B) A similar mechanical conflict arises at the organismal scale as longhorn crazy ants cooperatively haul a dead insect in the wild. Ant illustration adapted from *The Anteater’s Culinary Guide* by Ofer Feinerman and Ehud Fonio, with artwork by Sarit Bernard. (C) Approximate characteristic length scales of kinesin motors, ants and the robotic platform. (D) Robophysical realization of adversarial cooperation, in which two matched teams of wheeled robots pull a common cargo in opposite directions through compliant spring linkages. A reduced one-dimensional model and the corresponding minimal nondimensional equations for the mechanical and phase dynamics are shown below. (E) An individual wheeled robot and its principal components, including the compliant spring connection, force sensor, tracking markers, onboard controller and motorized wheel. (F) Sensing and control architecture for an individual robot, in which oscillator phase generates the motor command, the robot-cargo interaction returns local sensory feedback and a pre-trained phase-dependent predictor provides the reference for local phase adaptation.

Cooperative transport, therefore, provides a direct setting for asking how locally applied forces give rise to a coherent transport direction. Yet coherent motion of the cargo does not readily imply that every agent acts in the same direction at every moment. At the intracellular scale, a literal tug-of-war arises when kinesin and dynein remain engaged on the same cargo. Because their stepping and detachment depend on load, fluctuations in the engaged motor teams can shift dominance from one side to the other, producing pauses, reversals, and bidirectional runs [2, 3, 8]. At the scale of animal groups, longhorn crazy ants achieve directed transport through a changing division of labor. Ants at the leading edge pull, ants at the trailing edge lift to reduce friction, newly attached informed ants steer the group only transiently, and external constraints can drive collective oscillations that assist obstacle negotiation [6, 9, 10].

Across scales, these systems share a common form of mechanical feedback in which the shared load transmits force, and its motion changes what each active unit subsequently experiences (Fig. 1C). When the coupling is compliant, it can also store and release mechanical energy. Opposition can therefore produce stalling and force cancellation, but also load sharing, reversals, oscillations, or changes in which side dominates. This raises the possibility that, under intermittent pulling, the order of opposing actions may matter as much as their magnitude, and that the resulting loads may provide each agent with a local record of the interaction. Morphology and mechanical coupling have previously been used as embodied intelligence [11, 12] to simplify sensing, coordination, and control in active systems [13–19]. Whether such embodied mechanical feedback can organize persistent conflict into useful collective transport remains unclear.

Here, we formally explore adversarial cooperation in a robophysical tug-of-war, using two matched teams of spring-coupled wheeled robots that pull a shared cargo in opposite directions while coordinating their actions in time (Fig. 1D). Each robot combines a compliant spring connection, local mechanical sensing and onboard control (Fig. 1E), while its individual control architecture closes the loop between oscillator phase, motor command and sensory feedback without explicit inter-robot communication (Fig. 1F). The experiments reveal a form of embodied timing, in which relative pulling phase acts through the shared mechanics to select a transport direction, while the resulting loads provide local feedback for subsequent phase adaptation.

To uncover the origin of this effect, we develop a mechanically explicit model of intermittently pulling agents coupled through a compliant rope. Simulations reproduce the direction and approximate mean drift observed experimentally, while linear-response theory shows how compliant mechanics and ground resistance link relative pulling phase to a cycle-averaged power imbalance and, through the mean force balance, to directed transport. When the first phase harmonic dominates, the transport response is approximately sinusoidal, with extrema near quarter-cycle offsets. The resulting winning and losing states also produce distinct motor-current signatures, providing a local record of the mechanical interaction. We use these signatures to formulate a decentralized phase-adaptation rule that allows transient leadership to emerge and switch between teams as the collective transports high-friction cargo and navigates around obstacles.

## RESULTS

### Relative timing breaks the symmetry of mechanically matched tug-of-war teams

We set up a one-dimensional tug-of-war in which two mechanically matched teams of wheeled robots pulled against one another through compliant spring linkages (Fig. 2A). At the beginning of each trial, the individual oscillator phases were randomized, so that no inter-team phase difference was assigned initially. We represent the half-wave pulling gait in the mechanical model as

**FIG. 2.**
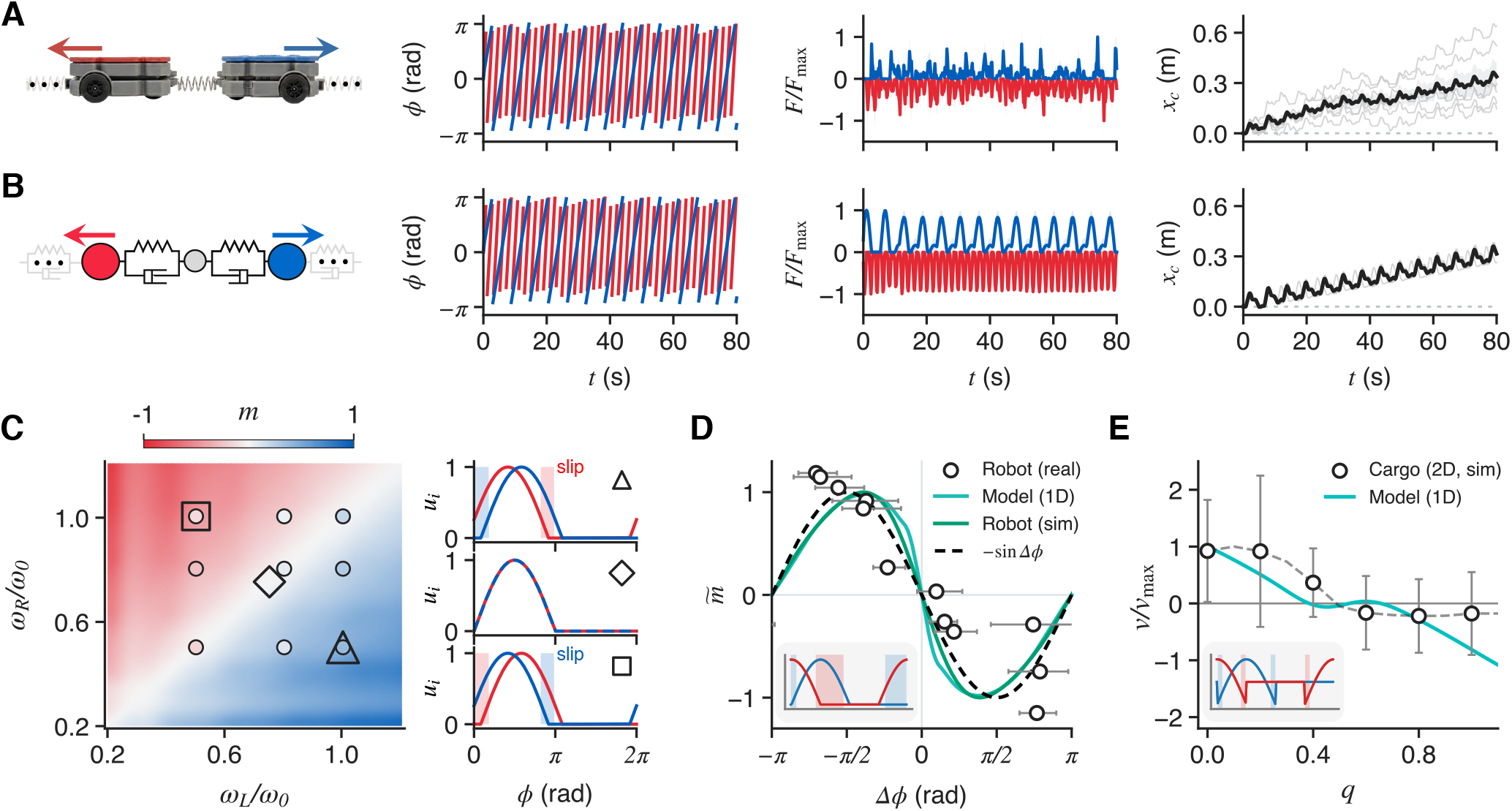
Relative timing breaks the symmetry of mechanically matched tug-of-war teams. (A) One-dimensional robophysical tug-of-war in which two red and two blue wheeled robots pull in opposite directions through compliant spring linkages. Representative wrapped phases *ϕ*_*i*_, normalized signed team pulling forces *F*_*i*_, and center displacement *x*_*c*_ are shown. Grey curves denote individual experimental realizations and the black curve denotes their mean. (B) Schematic of the reduced mechanical model, in which teams of two opposing active agents are coupled through Kelvin-Voigt elements. The model reproduces the phase evolution, opposing force cycles, and directional drift observed experimentally. For the representative experimental and simulation traces shown, the nominal intrinsic frequencies were *ω*_*L*_ = 3.1 rad s^*−*1^ and *ω*_*R*_ = 1.3 rad s^*−*1^. The force panel shows collective prescribed pull. In A and B, each team mean is normalized by its own peak magnitude. The model reproduces the direction and approximate mean displacement while underestimating variation between trials. (C) Signed transport order parameter *m* versus normalized team frequencies *ω*_*L*_*/ω*_0_ and *ω*_*R*_*/ω*_0_. The model frequency grid contains 51 *×*51 points, with each frequency pair averaged over a 16 *×*16 grid of starting-phase pairs. Circles show experimental condition means from 118 trials across nine conditions. Positive and negative values indicate right/blue and left/red dominance, respectively. The square, diamond, and triangle mark intrinsic-frequency pairs used in separate left-lagging, synchronous, and right-lagging controls. The adjacent plots show their pulling activations *u*_*i*_ versus common cycle phase; colored bands mark intervals during which one team is inactive while the other remains active. Frequencies in the heatmap are normalized by the robot’s motor drive, *ω*_0_ = 2*π ×*9.5*/*60 rad s^*−*1^. (D) Normalized transport order 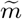 versus relative phase Δ*ϕ* = Φ_*R*_*−* Φ_*L*_. Open circles show 12 experimental trials at nominal frequency of 1 rad s^*−*1^; horizontal bars show the 5th–95th percentile temporal phase spread. The light green curve is the reduced 1D model response at 1 rad s^*−*1^, the dark green curve shows the robot simulation, and the dashed curve is the reference *−*sin Δ*ϕ*. Experimental and 1D model responses are normalized separately by positive, zero-offset fitted sine amplitudes. (E) Reduced-model prediction and two-dimensional cargo-simulation results under pull-and-brake forcing at 2 rad s^*−*1^ and Δ*ϕ* = *−π/*2. The normalized braking strength is *q* = *F*_hold_*/F*_0_ for the 1D model, which prescribes an outward holding force during inactivity, and *q* = *τ*_0_*/τ*_max_ for the cargo simulation, which limits the inactive-phase motor torque under the continuing wheel-speed controller. For each plot, the velocity is normalized by its own maximum magnitude over the displayed braking range. Positive and negative velocities favor the lagger and leader, respectively. Insets in D and E illustrate the pull-and-free (left) and pull-and-brake (right) commands. The error bars on the cargo-simulation scatter points denote 95% confidence intervals. Model parameters, detailed analysis, and two-dimensional simulations (Figs. S1, S6, and S9) are shown in Methods and the Supplementary Information.

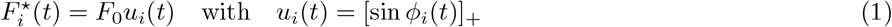

where [*z*]_+_ = max(0, *z*) and *F*_0_ is the common peak pulling force in the model. The direction of the force was determined by team membership, with the left and right teams pulling in opposite directions. The prescribed pull acts instantaneously and vanishes during inactivity. The model includes inertia, tension-only elastic coupling, baseline viscous and rolling resistance, and an activation-dependent resistance. The general chain model and its nondimensionalization are given in Methods and the Supplementary Information. For the comparison in Fig. 2B, four inertial masses are connected by Kelvin-Voigt elements. The recorded individual phases drive the numerical simulations, allowing us to compare the response with the experimentally observed timing, while the force panels separately compare normalized collective team-pulling forces. Both the experimental records and the simulations show mean motion towards the right (blue) team in the displayed, direction-aligned coordinates. The reduced one-dimensional model, along with the two-dimensional simulations shown in Supplementary Information (Fig. S7), reproduces the approximate mean displacement but underestimates the variation between robot experiment trials.

We quantify the net transport outcome by a signed macroscopic order parameter *m* = tanh(Δ*x*_*c*_*/δ*), where Δ*x*_*c*_ is the center displacement over the analysis window (*t*_obs_) and *δ* is a characteristic displacement scale. A positive value of *m* denotes right/blue dominance, whereas a negative value denotes left/red dominance. By varying the frequencies of the freely running teams, we observe left- and right-dominant regions separated by a near-neutral diagonal in the phase-averaged model map (Fig. 2C). The activation profiles alongside the heatmap display distinct phase-locked controls for the three marked intrinsic-frequency pairs. Coupling enables teams with unequal intrinsic frequencies to share a common realized frequency while maintaining a fixed relative phase once phase-locked. The triangle, diamond, and square markers indicate right-lagging, synchronous, and left-lagging pulling patterns, respectively. Their non-overlapping active intervals clarify which team remains active after the other becomes inactive, thereby establishing a fixed timing relationship.

At a fixed relative phase Δ*ϕ* = Φ_*R*_− Φ_*L*_, a negative Δ*ϕ* indicates that the right team lags behind. For identical mirrored teams, the settled transport response is odd and periodic in Δ*ϕ*. When the first phase harmonic is dominant, the mean velocity can be approximated as 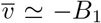 sin Δ*ϕ*, with *B*_1_ *>* 0 indicating a lagger-favoring regime. Over complete gait cycles, this results in *m* ≃ −tanh(*A* sin Δ*ϕ*), where *A*= *B*_1_*t*_obs_*/δ*. The sine approximation is valid for small accumulated displacements. The quarter-cycle extrema occur within the first-phase-harmonic approximation, though higher harmonics can shift these points. The experimental data, along with the reduced-model curve and the robot simulation curve displayed in Fig. 2D, show the corresponding dependence on the descending phase when normalized separately.

The transport mechanism follows from the mean force balance. Internal spring forces cancel each other out when summed across the assembly, while the cycle-averaged resistance related to activation does not necessarily cancel. The mechanical response links each robot’s velocity to its pulling activation, which allows for net drift even when the opposing mean pulls are equal and rolling resistance is linearized. Figs. S1 and S9 in the supplementary Information elaborate on this force balance, its relationship to cycle-averaged power, and the conditions that determine whether a participant has a lagger or leader advantage. This advantage can change if the inactive phase of the gait is modified. In the simplified model, when a pull-and-brake command is applied, the inactive half-cycle maintains a constant outward holding force. When there is a fixed quarter-cycle lag, increasing the holding force can reverse the overall transport direction, allowing the leader to gain an advantage (see Fig. 2E).

This reversal occurs without changing the imposed phase order. The reduced model further distinguishes off-phase holding from a constant outward force added throughout the cycle: on the same taut linear-elastic branch, with unchanged resistance activation, a constant offset changes the static extension but leaves the settled drift unchanged. Off-phase holding can therefore reverse transport through its temporal distribution, rather than simply through an increase in mean pulling force. In the two-dimensional robot simulations, additive torque offsets likewise retained the lagger advantage over the tested range (Fig. S11).

### Local current feedback organizes collective phase dynamics

The one-dimensional tug-of-war established that a quarter-cycle phase offset can select the winning team when the relative phase is imposed externally. We next asked whether a planar four-robot collective could recover this timing relation using only local mechanical feedback. Each robot was connected to one corner of a wheeled cargo through a linear spring (Fig. 1D) and carried its own phase variable *ϕ*_*i*_. The collective state was therefore determined by four phases, whereas robot *i* sensed only its own motor current *Ĩ*_*i*_, which provided a local proxy for the load transmitted through the springs and cargo.

To identify a local signature for each mechanical role, we first explicitly imposed the two quarter-period states during pre-training (Fig. 3A). The leading and delayed states produced reproducible yet distinct phase-dependent motor-current profiles. The current of the leading, losing robot exhibited a broad maximum, whereas that of the delayed, winning robot exhibited a sharper peak earlier in its active cycle (Fig. 3B,C). We represented the two profiles using separate radial-basis-function predictors for the winning and losing roles (Methods). The learned profiles reproduced the measured current traces with an RMSE of approximately 6.2%.

**FIG. 3.**
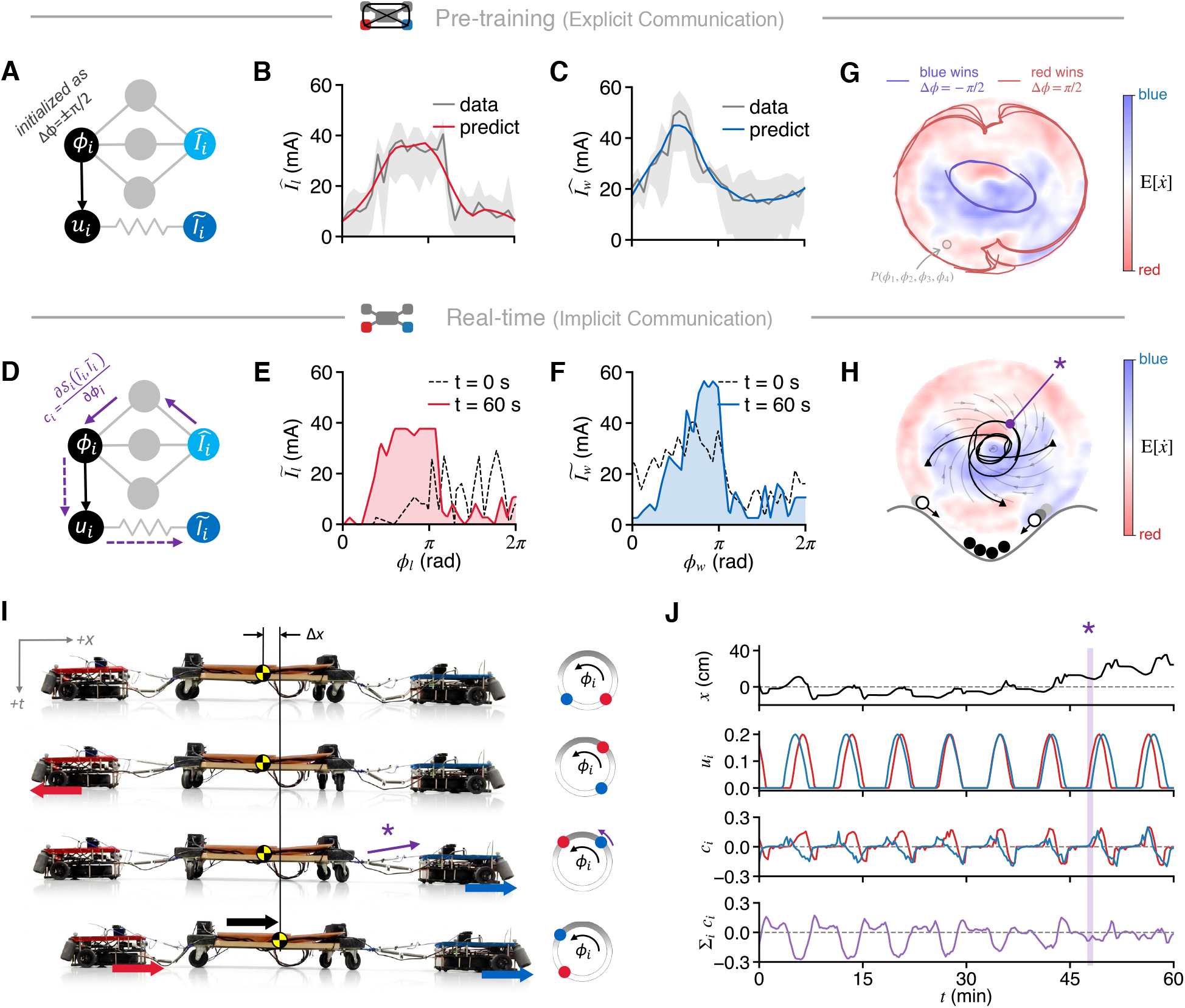
Learned current-phase relations enable self-organization of collective phase dynamics. (A) Pre-training architecture with an explicitly imposed quarter-period phase relation. The oscillator phase *ϕ*_*i*_ generates the motor command *u*_*i*_ and is mapped by a radial-basis-function predictor to a phase-dependent motor-current profile 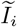. The compliant robot-cargo interaction produces the locally measured motor current *Ĩ*_*i*_. (B,C) Phase-resolved motor-current profiles measured for the losing (B) and winning (C) roles. Grey curves and bands show the measured profiles and their 95% confidence intervals; red and blue curves show the corresponding radial-basis-function predictions. The predictors reproduce the measured profiles with an RMSE of approximately 6.2% (*n* = 20 cycles). (D) Real-time architecture with implicit mechanical communication. Each robot compares its locally measured current with its assigned winning or losing current-phase profile, and the resulting local phase-adaptation signal *c*_*i*_ updates its oscillator phase. No robot receives the phases or currents of the other robots. (E,F) Motor-current profiles for robots assigned the losing (E) and winning (F) roles at the beginning of adaptation (dashed black curves) and after 60 s of local current-based adaptation (red and blue curves, respectively). Correlations with the assigned profiles increase from 0.22 to 0.64 for the losing role and from 0.82 to 0.91 for the winning role. (G) Two-dimensional latent representation of the four-robot phase state under explicit phase sharing. The background shows the conditional mean cargo velocity 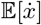, with blue and red regions corresponding to states that favor blue- and red-team motion, respectively. Blue and red trajectories show the explicitly imposed quarter-period states Δ*ϕ* = − *π/*2 and Δ*ϕ* = +*π/*2, and the open circle marks the initial collective state. (H) Representative latent trajectory under implicit current-based adaptation, overlaid on the same cargo-velocity map. Arrows indicate its temporal progression from the initial state towards a recurrent transport cycle. The schematic below illustrates the corresponding effective latent landscape, and the purple star identifies the state highlighted in (I,J). (I) Successive side-view snapshots and team phases during a representative pulling cycle. Both teams are initially inactive; the red team then becomes active and loads the compliant connections; the two teams subsequently overlap near maximum extension; and the red team enters its passive interval while the delayed blue team remains active, producing a rightward cargo displacement Δ*x*. (J) Cargo position *x*, red- and blue-team motor commands *u*_*i*_, individual phase-adaptation signals *c*_*i*_, and their sum Σ_*i*_ *c*_*i*_ during the same trial. The shaded interval corresponds to the starred state in (H,I). As the collective approaches a repeatable phase relation, the local phase corrections balance and their sum fluctuates around zero while the cargo advances towards the blue side. Two-dimensional simulations also recover this self-organized collective phase dynamics, see Fig. S8 in the Supplementary Information.

During real-time transport, explicit phase information was removed. Each robot compared its instantaneous current with its assigned winning or losing current-phase profile and used the resulting local phase correction to advance or delay its oscillator to increase agreement with that profile (Fig. 3D; Methods and Fig. S8 in the Supplementary Information). No robot received the phases or currents of the other robots. Nevertheless, the measured current profiles moved towards their respective targets during adaptation. In the representative traces shown, the phase-profile correlations increased from 0.22 to 0.64 for the losing role and from 0.82 to 0.91 for the winning role (Fig. 3E,F).

To visualize how these four independent phase updates reorganized the collective, we embedded the joint four-phase state into two dimensions using an autoencoder that operates on the sine and cosine of each oscillator’s phase (Methods). The latent representation was used only to analyze the collective dynamics and was not available to the robot controllers. Coloring this space by the observed cargo velocity revealed distinct regions associated with red- and blue-team dominance. Under explicit phase sharing, the two imposed quarter-period states traced reproducible reference trajectories through the corresponding regions of collective phase space (Fig. 3G).

Local current-based adaptation recovered the same direction-dependent organization without access to this global representation. A representative trajectory progressed from its initial state towards a recurrent orbit associated with the assigned winning team (Fig. 3H). The target direction was specified by assigning the winning and losing current profiles, but neither the relative phase nor the trajectory through collective phase space was prescribed. Both emerged through repeated interactions among each robot’s oscillator phase, its locally measured mechanical load, and the compliant cargo.

The physical cycle underlying this latent trajectory is shown in Fig. 3I. Both teams are initially inactive. The leading red team then pulls and stores energy in the springs, after which the two teams overlap near maximum spring extension. The delayed blue team remains active as the red team enters its passive interval, releasing the stored energy and advancing the cargo towards the blue side. Spring deformation simultaneously changes the current sensed by each robot, closing the local phase-adaptation loop. As the collective approaches a repeatable cycle, the individual corrections balance and their sum fluctuates around zero (Fig. 3J).

Thus, the decentralized controller does not encode an explicit global phase relation or a centralized transport plan. Instead, it uses learned phase-dependent current profiles associated with distinct mechanical roles. The compliant cargo couples the resulting local phase adaptations, turning them into global self-organization in collective phase space.

### Adversarial cooperation enables high-friction cargo transport

We tested whether the coordination produced by local motor-current feedback improved cargo transport under high friction. Four robots were coupled by springs to a common cargo and assigned a target along the positive *x* direction. We compared two controllers using the same physical platform. In the leader/follower baseline, one team was commanded towards the target while the other aligned its motion with the force transmitted through the cargo [20, 21]. Under adversarial cooperation, both teams remained active in opposing directions and adapted their phases using the learned current-phase profiles. We challenged both collectives with a cargo that had a high static-friction threshold of *f*_*s*_ = 10 N, exceeding the sustained pulling capacity of either team acting alone (Fig. S5 in the Supplementary Information).

Under leader-follower control, the cargo remained close to its initial position after one hour. By contrast, cooperation emerging from our embodied, adversarial collective transported the same cargo towards the target over the same interval (Fig. 4A,B). The difference was not simply a transition from slow to uniformly faster motion. Rather, the leader-follower controller occupied a narrow and strongly asymmetric trajectory in the collective phase embedding, alternating between states in which the leader advanced and states in which it slipped backward (Fig. 4C). Embodied adversarial cooperation produced a broader, approximately symmetric distribution concentrated closer to the center of the latent space (Fig. 4D). Forward transport, therefore, accumulated as a small directional bias over repeated cycles of advance and recoil rather than as steady translation.

**FIG. 4.**
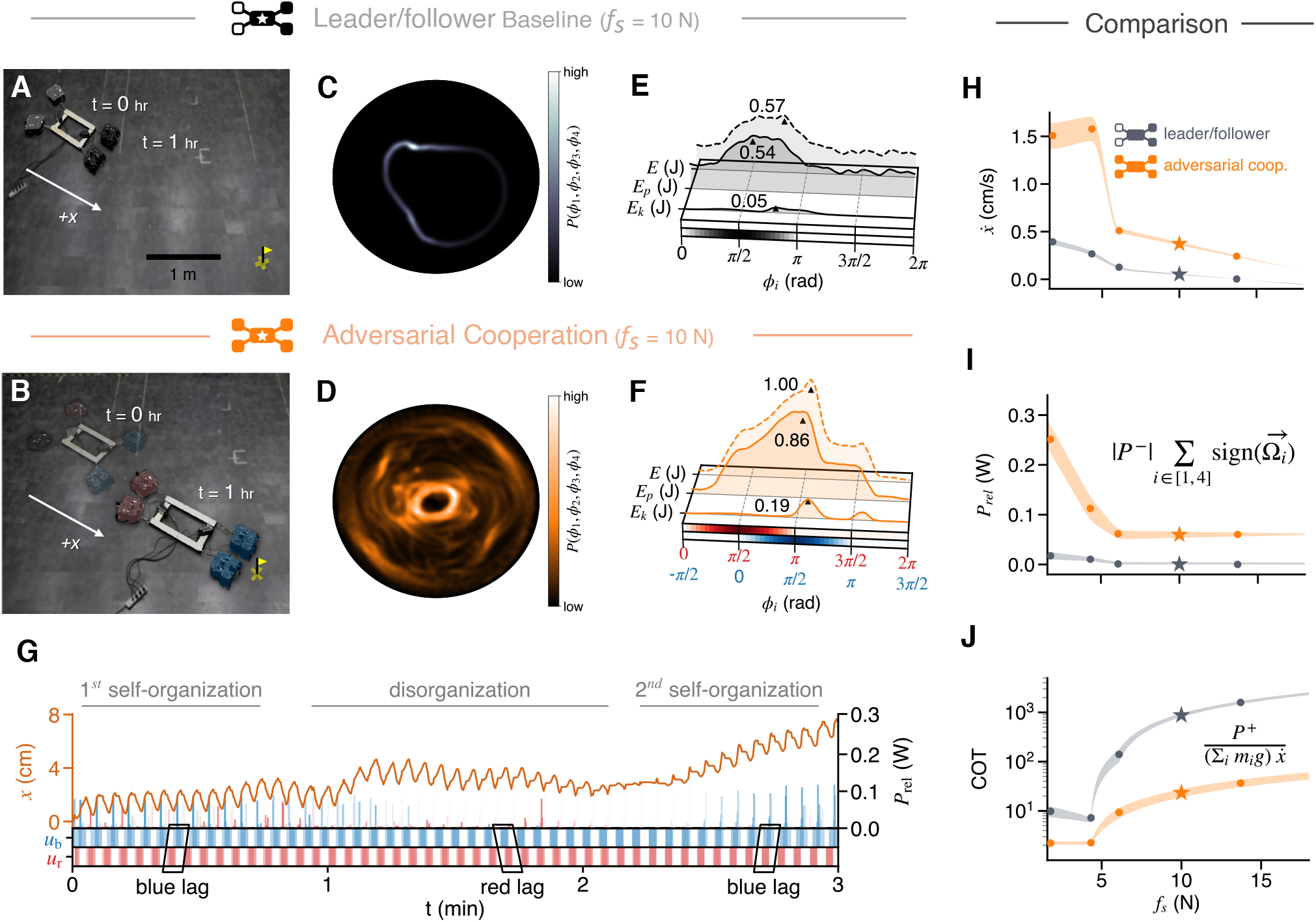
Embodied adversarial coordination enables high-friction cargo transport through phase-organized energy storage and release. (A,B) Transport of a cargo with static-friction threshold *f*_*s*_ = 10 N towards the target marked in yellow. Under (A) leader-follower control, one team is commanded towards the target while the second team follows the mechanically transmitted pull; the cargo remains near its initial position after one hour. Under (B) adversarial cooperation, both teams remain active in opposing directions and adapt their phases through local motor-current feedback; the same cargo is transported towards the target. (C,D) Probability densities of collective trajectories in the two-dimensional phase embedding under (C) leader-follower control and (D) adversarial cooperation. Brighter regions denote more frequently occupied states, and the open circle marks the initial state. (E,F) Phase-aligned total, potential and kinetic mechanical energies under (E) leader-follower control and (F) adversarial cooperation. In (F) the red and blue phase axes show the activation phases of the leading and delayed teams, respectively. Curves are averaged over *n≥* 100 cycles (RMSE_*ϕ*_ = 0.23 rad). (G) Representative adversarial-transport trial showing cargo position *x*, energy-release rate *P*_rel_, and red- and blue-team commands *u*_*r*_ and *u*_*b*_. The collective undergoes an initial self-organization, a temporary loss of phase organization and a second self-organization into the blue-lag state. (H-J) Mean cargo speed 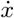, energy-release rate *P*_rel_ and cost of transport, respectively, as functions of cargo friction for leader-follower control (grey) and adversarial cooperation (orange) (also see Figs. S5 and S6 in the Supplementary Information). Stars identify the *f*_*s*_ = 10 N condition shown in (A-G). Shaded bands denote 95% confidence intervals; *n ≥*100 cycles per condition.

To identify the mechanical origin of this bias, we aligned individual cycles by the activation phase of the robots and measured the potential, kinetic and total mechanical energy of the system (Fig. 4E,F). The adversarial collective accessed a substantially larger energy range. The peak total, potential and kinetic energies shown in the figure were approximately 1.00, 0.86 and 0.19 J, respectively, compared with 0.57, 0.54 and 0.05 J under leader-follower control.

More importantly, the timing of energy storage and release differed between the two controllers. Under leader/follower cooperation, the energy rose and fell around the activation of a single leading team. Under adversarial cooperation, the leading red team first loaded the compliant connections to store energy, while the delayed blue team remained active as the stored elastic energy was released. The delayed team could therefore retain traction during energy release, converting the mechanically opposed pulling cycle into displacement towards the target.

The high-friction load also perturbed this organization dynamically. During the first minute of the representative trial, the collective organized into a state in which the blue team lagged and the cargo advanced towards the target (Fig. 4G). The phase relation was temporarily disrupted between approximately one and two minutes, shifting the largest energy-release events towards red-team activation and briefly reversing the cargo motion. This reversal altered the locally measured current profiles and reactivated phase adaptation. A second self-organization event then restored the blue-lag state and renewed forward transport. The controller, therefore, did not maintain a rigidly prescribed phase offset; instead, it repeatedly recovered a transport-producing state through continued interaction with the load (see Fig. S6 in the Supplementary Information).

We next varied the cargo friction from 1 to 15 N. For *f*_*s*_ ≤5 N, adversarial cooperation transported the cargo approximately three to four times faster than leader/follower control, while producing greater phase-localized energy release and a lower cost of transport (Fig. 4H-J and Fig. S5 in the Supplementary Information). The speed of both controllers decreased sharply as the load approached the frictional limit, but the adversarial collective retained an advantage in both speed and energy release under near-critical conditions. Persistent opposition, therefore, did not merely coexist with transport. By increasing the mechanical energy stored in the compliant system and timing its release to coincide with the activation of the delayed team, adversarial coordination extended the range of loads the collective could move.

The phase trajectories nevertheless became broader and less regular as cargo friction increased. This raises a further question: whether such repeated loss and recovery of the organized state is merely a cost of operating near the transport limit, or whether it can itself support adaptive collective behavior.

### Environment-mediated role switching enables obstacle navigation

The high-friction transport experiments showed that external loading could temporarily disrupt the organized phase relation before the collective recovered it. We next asked whether these departures from the nominal winning state could become useful in a geometrically constrained environment. We placed the four-robot collective in a maze with two orthogonal turns, with an intermediate target at each turn to provide high-level route guidance (Fig. 5A).

**FIG. 5.**
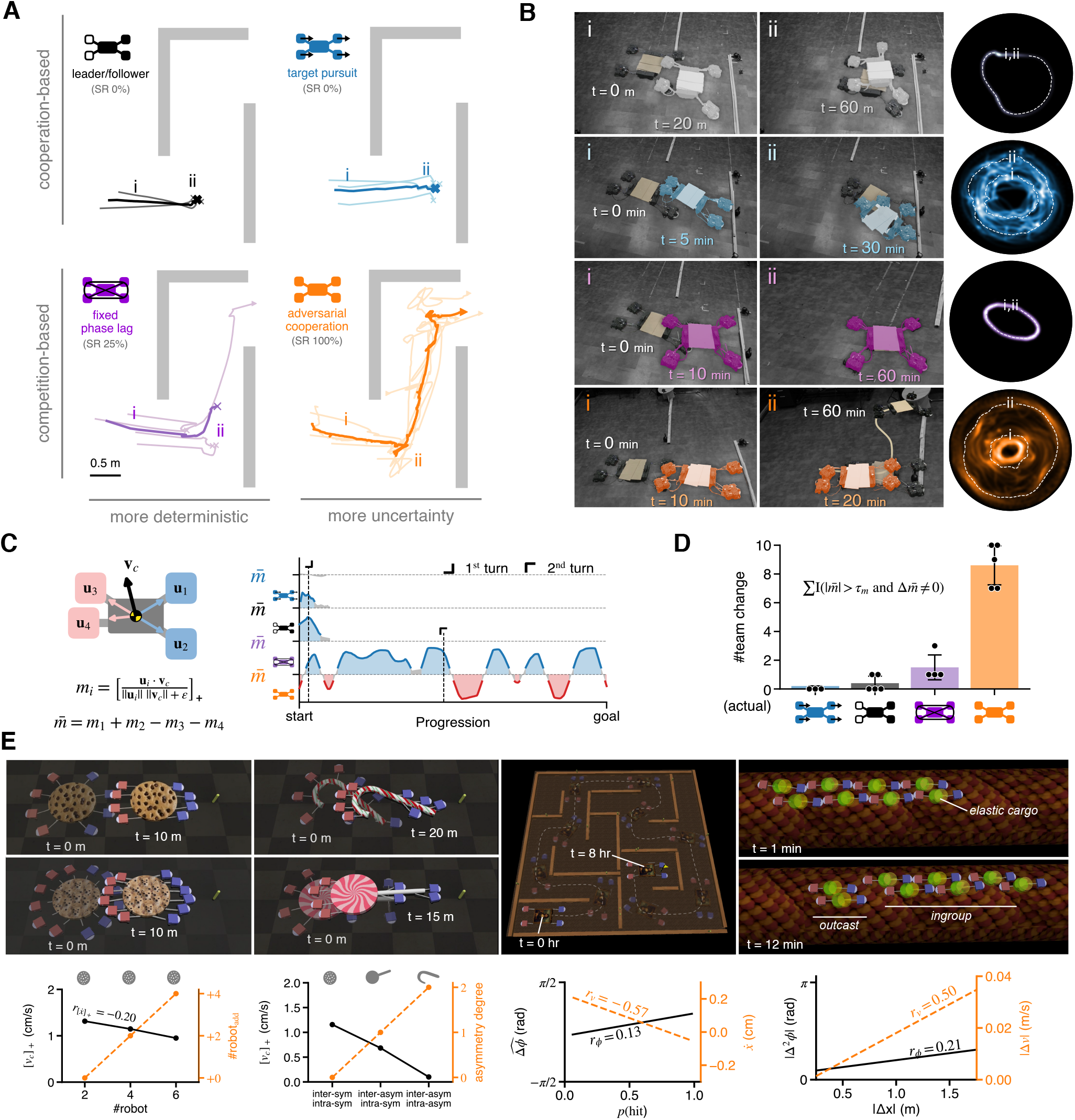
Environment-mediated role switching enables obstacle navigation. (A) Cargo trajectories through a maze containing two orthogonal turns under four coordination schemes. Leader-follower control (black) assigns one team to move towards the target while the second follows the mechanically transmitted pull. Target pursuit (blue) commands each robot independently towards its assigned target. Fixed phase lag (purple) uses explicit communication to enforce a constant quarter-period phase offset and a predetermined winning team. Adversarial cooperation (orange) uses local motor-current feedback without explicit inter-robot communication. Labels give the observed success rate for each scheme; (i) and (ii) mark states before and after the first wall encounter. (B) Experimental snapshots corresponding to states (i) and (ii) in (A), together with the associated probability densities in the two-dimensional collective phase embedding. Brighter regions denote more frequently occupied states. (C) Definition of the individual motion-alignment measure *m*_*i*_ = [***u***_*i*_ *·* **v**_*c*_*/*(∥****u***_*i*_*∥∥**v**_*c*_∥ + *ϵ*)]_+_. The unsigned sum _*Σ i*_ *m*_*i*_ measures collective alignment, whereas the signed difference 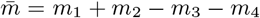 identifies the mechanically dominant team. The plots on the right show the evolution of 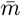 along the route for each coordination scheme. Positive values denote blue-team dominance and negative values denote red-team dominance. (D) Number of effective winning-team changes per trial. A role switch was counted when 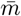 crossed the 25% dead band around zero and subsequently entered the opposite team-dominant state. Points denote individual trials, bars show trial means and error bars indicate 95% confidence intervals; *n ≥*4 trials per condition. (E) Additional demonstrations across changes in robot number, cargo geometry, maze structure, and simulated environments (also see Figs. S8–S14 in the Supplementary Information).

We compared four coordination schemes. In the leader-follower controller, one team moved towards the target while the other followed the force transmitted through the cargo [20, 21]. In target pursuit, each robot independently moved towards the target using its own state space [22–24]. The fixed-lag controller used explicit phase communication to maintain a prescribed quarter-period offset and a predetermined winning team (ablation). Adversarial cooperation retained the same nominal winning and losing assignments, but each robot adapted its phase using only its locally measured motor current (proposed).

Both cooperation-based controllers failed to negotiate the first turn in all reported trials. Under leader-follower control, a leading robot became wedged against the wall, whereas under target pursuit, a pushing robot lost effective contact and displaced the cargo away from the route. Fixed-lag control succeeded in one of four trials; in the remaining trials, maintaining the predetermined winning team caused the collective to stall at the first turn. By contrast, adversarial cooperation successfully traversed both turns in every reported trial, despite the absence of explicit inter-robot communication (Fig. 5A,B).

The corresponding latent-state distributions reveal how the four controllers responded to wall contact. Leader-follower and fixed-lag control remained confined to comparatively narrow trajectory families after becoming trapped, whereas target pursuit spread across a broader but unproductive region of collective phase space. Adversarial co-operation instead occupied two recurrent radial regions: one associated with forward transport towards the target and another reached after wall contact, when the cargo moved backward or sideways (Fig. 5B). Obstacle encounters, therefore, did not merely add noise to an otherwise fixed phase cycle; they drove transitions between distinct collective transport states.

To quantify these transitions without relying on the latent representation, we measured how strongly each robot’s commanded motion aligned with the instantaneous cargo velocity and combined these contributions into a signed team-dominance measure 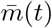 (Fig. 5C; Methods). Positive values of 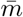 denote blue-team dominance, whereas negative values denote red-team dominance. Although the blue team remained the nominal winner in the fixed-lag and adversarial controllers, the realized mechanical roles depended on both the controller and the environment. Leader-follower control showed no sustained reversal before stalling, target pursuit showed no coherent winning team, and fixed-lag control remained largely dominated by its predetermined winner. Under adversarial cooperation, by contrast, 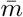 repeatedly changed sign.

We counted a role switch only when 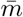 crossed the 25% dead band around zero and subsequently entered the opposite team-dominant state. Adversarial cooperation produced substantially more such switches than any of the three comparison controllers (Fig. 5D). These reversals were mechanically triggered. When a wall stopped a blue-team robot, its spring could no longer extend, and the resulting drop in motor load changed the local current-phase relation on which phase adaptation was based. The nominally losing red team could then become transiently dominant and pull the cargo backward or sideways away from the obstruction. Once the collective cleared the wall, the load signals changed again, and the robots reorganized into a blue-dominant state directed towards the next target. The route was prescribed, but the sequence of effective leaders required to follow it was selected in real time through interaction among the robots, the compliant cargo, and the environment.

Additional demonstrations across changes in robot number, cargo shape, attachment configuration, maze structure, and simulated environment suggest that this mechanism is not tied to a single collective geometry, and that different interaction types shape collective function in distinct ways (Fig. 5E and Figs. S8–S14 in the Supplementary Information). Specifically, increasing the number of robots beyond pre-training conditions caused a proportional decline in transport speed, though forward velocity remained strictly positive. A similar trend emerged under asymmetry: when cargo geometry increased the distance from the contact points to the center of mass for the opposing team (red), it impeded full heading alignment, reducing transport speed while maintaining positive overall movement. In the complex maze, correlation plots showed that shifts from lagging to leading were associated with temporary changes from forward to backward cargo motion following wall impact. Finally, spatial separation between collective sub-units, measured by the inter-unit distance |Δ*x*|, impeded phase-difference alignment across the system, disrupting transport-speed synchronization and ultimately inducing group separation.

Role switching allowed the collective to escape local mechanical traps. Contact with obstacles changed the loads sensed by individual robots, and the resulting phase adjustments reorganized the pulling sequence so that the cargo could move away from the obstruction.

## DISCUSSION

Across living matter, competing or alternating stresses often provide the route by which a system changes state. Repeated stiffening and fluidization can ratchet an incomplete contractile band through cytokinesis; time-dependent interfacial tension and differential migration can drive T1 neighbor exchanges; competition between cellular polarities and active boundaries can select coherent epithelial rotation; and distributed active stresses can organize tissue-scale defects and architecture [25–29]. These studies show how opposition and temporal ordering generate shape, topology, and flow. Our work asks a different question: when groups of agents act through the same compliant body, can timing alone determine which side gains the advantage? By isolating this contest, we identify relative phase as a key control variable and show how compliant dynamics and gait-dependent resistance combine to produce net transport.

Our results identify relative timing as a control variable for mechanical dominance, whose effect depends on the mechanics and actuation of the coupled system. Previous work has shown that elastic and viscoelastic coupling can select collective modes and spatiotemporal order, that morphology and collisions can add a mechanical layer to decentralized learning, that nonlinear elasticity can rectify nearly diffusive actuation into directed navigation, and that noisy active particles can serve as physical computational reservoirs [13, 16, 30–33]. Our work extends this view to internal conflict and reveals a dual role for mechanics. The same deformation field that redistributes mechanical work also generates local current-phase fingerprints of global mechanical roles. The body, therefore, does not merely compute; it adjudicates a contest.

The braking results clarify why timing must be understood together with the mechanical response. In the reduced model and robot simulations, changing the inactive-phase actuation can reverse the winning team without changing the imposed phase order. A lagging team can therefore win or lose depending on how the actuators interact with the load throughout the cycle. The placement experiments provide a complementary test: same-side teams moved faster with in-phase pulling, whereas opposing teams benefited from a phase offset (Fig. S9). Together, these results show that the advantageous timing pattern depends on both the mechanical arrangement and the actuation protocol. Local load feedback makes this dependence useful for adaptation. Environmental contact changes the loads experienced by individual robots, modifying their phase corrections and allowing the collective to reorganize its timing. This provides a second route to changing mechanical dominance: braking reverses the advantage at fixed phase order, whereas decentralized feedback allows the phase relationship itself to evolve. The shared mechanics thus link the selection of a transport direction to the local signals that can revise that direction when the collective encounters an obstacle.

This embodied mechanics-based view also changes how transient leadership can be interpreted. Alternating temporal leaders guide intermittent collective motion in sheep; conspicuous changes in body orientation create effective leaders during fish U-turns; leadership is distributed across distinct functions in octopus-fish hunting groups; and phase-leading, phase-lagging, and shared leadership patterns emerge in synchronized human networks [34–37]. These studies generally infer leadership from trajectories, social influence, or information-flow networks. Here, the causal chain can be closed: environmental contact changes motor load, motor load changes local phase adaptation, and phase adaptation changes which team dominates transport. A role switch, therefore, requires neither a new global instruction nor an explicit role-switching policy. Mechanical dominance is not a permanent property of any agent or team, but a dynamical state of the coupled agent-body-environment system. In this sense, the environment does not simply perturb the computation performed by the collective; it becomes one of its physical inputs. This interpretation complements recent evidence that human and animal groups can escape established coordination states by reorganizing phase, coupling, or leadership relationships rather than by preserving a fixed hierarchy [38, 39]. In rival Siamese fighting fish (*Betta Splendens*), repulsive interactions can lead to behavioral synchronization as the fight drives behavioral and even brain-transcriptomic synchronization between the two opponents [40].

The robophysical tug-of-war provides a minimal system in which phase, frequency, compliance, friction, geometry, and local sensing can be varied separately, while theory, simulation, and hardware share the same signed observable. This tractability yields predictions that can be sought beyond the present platform: exchanging the opposing groups should reverse the response; changing stiffness or forcing frequency can shift or reverse the advantageous lead-lag branch; and changes in local load should precede changes in effective mechanical leadership. Reconfigurable swarms and morphology-assisted robots have shown how changing body organization, connection mode, or contact mechanics can broaden collective behavior [41–44]. Most robotic designs avoid unresolved competition between actuators. Their force or velocity directions are generally aligned to reinforce a prescribed motion; when actuators are arranged in opposition, their cancellation is typically used to suppress one mode and isolate another, as when oppositely directed side-fin actuation cancels net translation to produce in-place rotation [45]. Collective transport controllers similarly align individual movement or force vectors towards a common direction [21, 46].

Here, by contrast, we deliberately retain mechanical opposition. By organizing competing actuation directions in time, the collective converts what would otherwise be force cancellation into transient mechanical advantage and effective role switching. Here, functional reorganization occurs without changing morphology or replacing the individual control policy; the collective instead moves between mechanically distinct role configurations. The present robots still receive prescribed targets and pre-trained role profiles, and whether this mechanism scales to larger, heterogeneous or dynamically connected groups remains an open question. Even so, the broader principle, perhaps, extends beyond this implementation. A collective need not eliminate disagreement before it acts; rather it can use a shared body to rectify that disagreement into useful work.

## METHODS

**One-dimensional mechanical model**

The one-dimensional compliant rope consists of *N* nodes, with lumped mass *m*, and active agents with mass *M* (*M*≫ *m*). Adjacent rope nodes are connected by tension-only Kelvin-Voigt elements with rest spacing *l*_0_, axial stiffness *EA/l*_0_ and internal damping coefficient *γ*_*r*_. The tension in the span connecting nodes *i* and *i* + 1 is given by

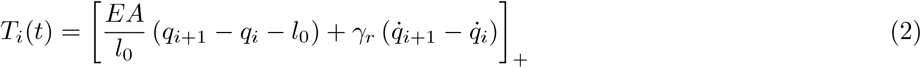

where [*z*]_+_ = max(0, *z*) enforces the tension-only constraint. The equation of motion for an interior node after summing the mechanical and active forces is given by 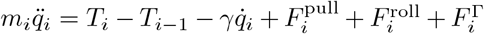, with the corresponding one-sided forms used at the two free endpoints. An agent *j* occupying a node *p*_*j*_ can belong to either the left or right team, represented by *s*_*j*_ = −1 or *s*_*j*_ = +1, respectively. Its phase generates the half-wave activation *u*_*j*_(*t*) = [sin *ϕ*_*j*_(*t*)]_+_. The node-wise pulling force and activation-dependent resistance are then given by, 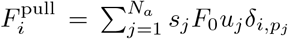 and 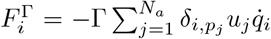. During inactivity, the baseline pull and activation-dependent resistance vanish. The agent nodes additionally experience a regularized rolling-resistance force

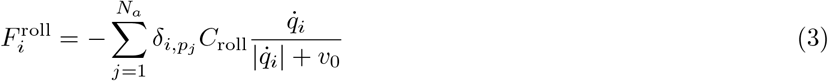

Within each team, oscillator phases evolve through their intrinsic frequencies, Kuramoto coupling and phase noise:

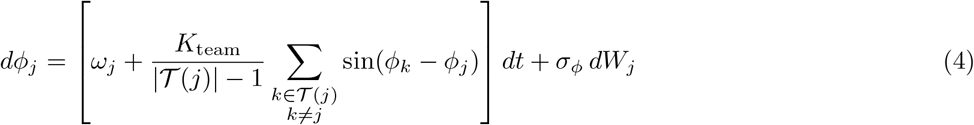

Here, *T* (*j*) denotes the team containing agent *j*. The complete mechanics derivation, free-end equations, and prescribed-phase protocols are given in the Supplementary Information.

### Nondimensionalization and numerical simulations

We use the rope-node spacing *l*_0_ as the length scale, *EA* as the force scale and 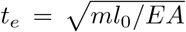 as the elastic timescale. Dimensionless position and time are defined as *x*_*i*_ = *q*_*i*_*/l*_0_ and *τ* = *t/t*_*e*_. After scaling force by *EA*, the dimensionless equation for an interior node becomes

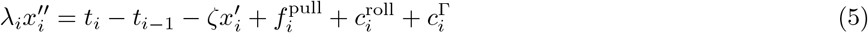

where primes denote derivatives with respect to *τ*,*λ*_*i*_ = *m /m*, and 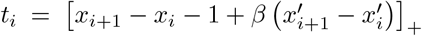 . The dimensionless damping parameters are 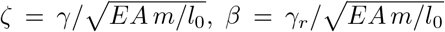, and 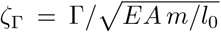. Rolling resistance, intrinsic frequencies, intra-team coupling, and phase noise are rescaled using the same force and time scales. Their definitions and the conversion to the numerical implementation are provided in the Supplementary Information, Tables 2, 3, and 5.

The Fig. 2 calculations use four agent nodes connected by three tension-only springs with no cargo. All calculations retain nonlinear rolling resistance and instantaneous activation-dependent resistance. Panel B uses recorded individual phase histories, measured initial positions, and initial velocities. Its integration step is 10^*−*3^ s, with output every 0.2 s.

Panel C prescribes freely running team phases on a frequency grid spanning 0.2-1.2 rad s^*−*1^ with spacing 0.02 rad s^*−*1^. Each frequency pair is averaged over a uniform 16 ×16 grid of starting-phase pairs. Model displacement is measured over 30-60 s after relaxation while experimental condition means use 10-40 s. The plotted signed transport order parameter is the ensemble mean of the individual values tanh(Δ*x*_*c*_*/δ*).

### Linear-response analysis

We consider identical mirrored teams with synchronized agents, fixed relative phase, taut spans, and velocities sufficiently small to linearize rolling resistance. Let **x**^(0)^ denote the equilibrium under the common mean pulling force and **y** = **x** − **x**^(0)^. The mechanical response obeys

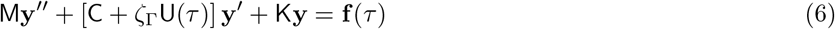

where **f** is the mean-subtracted applied force, M = diag(*λ*_*i*_), K = D^T^D, and C = *ζ*I + *β*D^T^D + (*σ*_roll_*/ε*_0_)P_*a*_. Here, D is the chain incidence matrix, P_*a*_ selects agent nodes, and 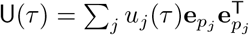. For a settled response with bounded relative coordinates, all nodes share the mean velocity *V* . Under pull-and-free forcing, summing and averaging the node equations gives *D*_tr_ *V* = −*ζ*_Γ_ *∑*_*j*_ ⟨*u*_*j*_ *v*_*j*_⟩_cyc_ with *D*_tr_ = *Nζ* + (*N*_*a*_ *σ*_roll_ */ε*_0_) *>* 0, where 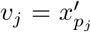. For the physical waveform *f*_0_*u*_*j*_, the right-minus-left supplied pulling-power difference is 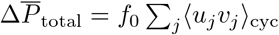. Consequently,

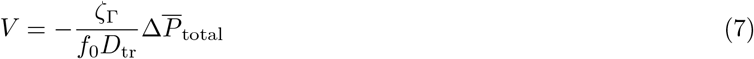

This proportionality applies to pull-and-free forcing with linearized rolling resistance. The full nonlinear rolling law and the additional holding work require the corresponding terms to be retained in the mean balance. We use Φ_*L*_ = Ω*τ* and Φ_*R*_ = Ω*τ* + Δ*ϕ*, so that Δ*ϕ <* 0 denotes the right team lagging. Reflection symmetry gives an odd, periodic phase response. Its leading phase harmonic yields the sine approximation, but its coefficient and the higher harmonics depend on frequency and mechanical parameters. The controlled small-force expansion, power accounting, and conditions for lagger advantage and braking-induced reversal are derived in the Supplementary Information.

### Phase-response and braking analysis

For Fig. 2D, we use 12 experimental trials with equal nominal frequencies of 1 rad s^*−*1^. For each team *α ∈*{ *L, R }*, the collective phase was calculated as the circular mean of the individual robot phases,

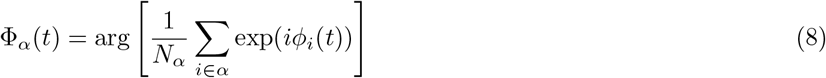

The trial-level phase offset is the circular mean of Δ*ϕ*(*t*) = Φ_*R*_(*t*) −Φ_*L*_(*t*) over 10-(10 + 8*π*) s. Horizontal bars show the 5th-95th percentile temporal phase spread around that mean. Points represent individual trials, without phase folding or binning. The model prescribes fixed phase differences over [− *π, π*] at 1 rad s^*−*1^. Mean velocity is calculated over the final four cycles of a 64-cycle integration, and the transport order parameter is 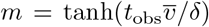, with *t*_obs_ = 8*π* s. Experimental and model responses are normalized separately. For each dataset, define *z*_*r*_ = −sin Δ*ϕ*_*r*_ and fit

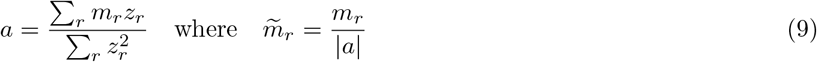

Both fitted amplitudes are positive for the displayed data. This normalization preserves the sign and compares the phase-dependent shape, rather than the absolute transport amplitude and the horizontal experimental bars describe temporal phase variation. For Fig. 2E, the frequency is 2 rad s^*−*1^ and Δ*ϕ* = − *π/*2. We vary *q* = *F*_hold_*/F*_0_ while retaining the original activation in the resistance term. Mean velocity over the final four cycles of a 64-cycle integration is normalized by its maximum positive value over 0≤ *q*≤ 1.2. Positive velocity favors the lagging right team and negative velocity the leading left team.

### Robotic platform and low-level control

The experiments used differential-drive wheeled robots coupled through compliant spring elements. The mass of each robot was 2.9 kg, while an additional 2.0 kg was attached in the transport experiments. The compliant elements had a lumped spring constant of 260 N m^*−*1^. The wheel radius and wheel separation were 0.033 m and 0.016 m, respectively. The controller was updated every Δ*t* = 0.1 s. Full physical parameters and robot-specific controller gains are listed in Supplementary Tables 2 and 3.

At discrete time *t*_*n*_ = *n*Δ*t*, robot *i* updates its internal phase according to *ϕ*_*i*_(*t*_*n*+1_) = *ϕ*_*i*_(*t*_*n*_)+*ω*_*i*_Δ*t*+*f*_*c,i*_(*t*_*n*_)+*ξ*_*i*_(*t*_*n*_), where *ω*_*i*_ is the intrinsic phase frequency, *f*_*c,i*_ is the coupling or adaptation signal and *ξ*_*i*_ is zero-mean Gaussian noise. The normalized pulling command is *u*_*i*_(*t*_*n*_) = [sin *ϕ*_*i*_(*t*_*n*_)]_+_ so that each robot alternated between active and passive half-cycles. In this setup, motor current provides a local proxy for mechanical load. To remove robot-specific no-load current, the measured current is corrected using a dead band:

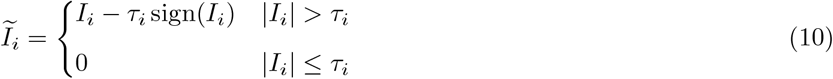

The current-feedback gains and dead bands were calibrated individually to account for differences in hardware response and static and kinetic friction. Finally, the commanded forward velocity combines encoder-based velocity feedback, phase-dependent actuation and current-based load compensation: 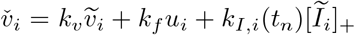. The current gain switches between near-static and finite-speed values according to whether *v*_*i*_ is below or above the velocity threshold *τ*_*w*_. Heading regulation is applied independently through 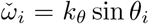 sin *θ*_*i*_, and the forward and angular commands are converted into differential left- and right-wheel duty cycles. The complete wheel-encoder conversion, gain-switching rule and differential-drive equations are given in the Supplementary Information.

### Phase coupling and learning of current-phase profiles

We use two phase-coupling protocols. In the explicit phase-sharing protocol, robots within the same team are coupled through 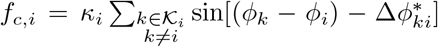, with coupling strength, *κ*_*i*_ *>* 0 and _*i*_, the set of indices of all robots coupled to robot *i*. This protocol approximates the phase-locked-team limit of the mechanical model.

For current-based decentralized protocol, we first collect pairwise tug-of-war calibration data with the coupling term set to 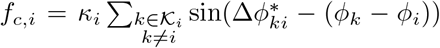, where the target phase difference 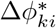 is set to −*π/*2 for a winning robot and 0 for a losing robot. We record each robot’s phase and dead-band-corrected current before fitting separate phase-dependent current profiles for winning and losing roles. Each profile is represented using five radial basis functions,

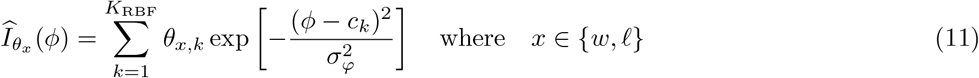

with *K*_RBF_ = 5 and *σ*_*Φ*_ = 1.0. The coefficients are obtained by minimizing the least-squares prediction error between the predicted and measured current. During decentralized trials, robot *i* evaluates the local phase-current alignment score 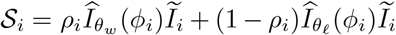 where *ρ*_*i*_ = 1 selects the winning profile and *ρ*_*i*_ = 0 selects the losing profile. Its phase-adaptation signal is

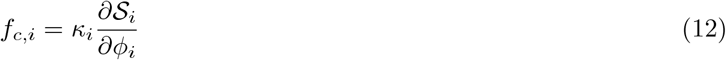

Each robot therefore adjusts its phase using only its own oscillator phase and motor current. No phases, current measurements, or adaptation signals are exchanged between robots. The phase derivative is evaluated while holding the measured current fixed, giving 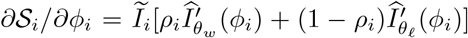. Environmental load changes therefore modulate the local phase corrections through the measured current. Differences between these corrections change the robots’ relative phase evolution; the resulting pulling order and transport direction are evaluated from their trajectories (Supplementary Information).

### Latent phase-space representation

For visualization of the collective phase dynamics, the four oscillator phases are embedded as the eight-dimensional sine-cosine vector **s**(*t*) = (sin *ϕ*_1_, cos *ϕ*_1_, …, sin *ϕ*_4_, cos *ϕ*_4_)^T^. A two-layer encoder with hyperbolic-tangent activations maps **s** to the two-dimensional latent state **z** = (*z*_1_, *z*_2_)^T^, and a corresponding two-layer decoder reconstructs the input: 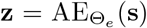 and 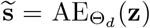. The autoencoder is trained using data from both mechanically coupled and decoupled experiments. Its objective combines mean-squared reconstruction error with a temporal contrastive term, ℒ_AE_ = ℒ _rec_ + *λ*_con_ ℒ_con_, with *λ*_con_ = 0.01 and contrastive temperature *α* = 0.1. The complete contrastive objective is given in the Supplementary Information.

Autoencoder accuracy is evaluated on a held-out test set using the reconstruction error

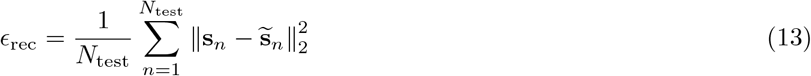

To generate continuous displayed trajectories while preserving their cyclic geometry, the latent coordinates are mapped to the unit sphere, smoothed using a second-order Savitzky-Golay filter and interpolated using cubic splines before being projected back onto the latent disk.

### Mechanical dominance and transient leadership

We define leadership by measuring the alignment between each robot’s commanded motion and the observed cargo velocity. Let **v**_*o*_(*t*) denote the windowed cargo center-of-mass velocity and let **ĥ**_*i*_(*t*) denote the heading unit vector of robot *i*. The commanded motion vector is ***u***_*i*_(*t*) = *u*_*i*_(*t*) **ĥ** _*i*_(*t*), and the individual contribution to cargo motion is

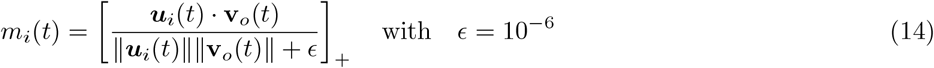

The unsigned sum Σ _*i*_ *m*_*i*_ quantifies the total alignment between the robot commands and cargo motion. Differential team dominance is defined as 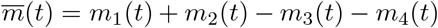, where robots 1 and 2 form the right/blue team and robots 3 and 4 form the left/red team. Positive values therefore denote right-team dominance and negative values denote left-team dominance. A team is classified as mechanically dominant only when 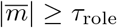, with *τ*_role_ = 0.25. A role switch is recorded when the trajectory leaves one team-dominant state, crosses the dead-band region and subsequently enters the opposite team-dominant state.

### Calibration of the reduced model

Robot mass and spring stiffness were fixed at *M* = 2.9 kg and *k*_*s*_ = 260 N m^*−*1^. An initial set of effective force, resistance, and rest-separation parameters was obtained by fitting all four robot-position trajectories in 52 experimental trials over 10.5-40 s, using measured individual phases as inputs. Winner labels and a prescribed phase-transport relationship were not used in the fitting objective. The preliminary fit allowed resistance activation to relax with a time constant *τ*_*r*_. Subsequent comparison with the six Figure 2A records motivated instantaneous activation, *τ*_*r*_ = 0, and a sensitivity calculation selected Γ = 379.89 N s m^*−*1^. The calibrated model uses symmetric teams, intra-team coupling strength *κ* = 0.50, dimensionless integration step 0.05 and physical output interval 0.05 s. The complete parameter values and trajectory comparisons are provided in the Supplementary Information.

## DATA AND CODE AVAILABILITY

Code, demonstrations, and data are available at https://github.com/Arthicha/TugOfCargoTransport.

## SUPPLEMENTARY MOVIES

Movie S1. Quarter-period phase lag in Tug-of-War (https://youtu.be/XQK2eLwMaYI)

Movie S2. Cooperative cargo transport in Tug-of-War (https://youtu.be/IBXKxI3KKZ8)

Movie S3. Adversarial cooperation enables high-friction cargo transport (https://youtu.be/wFvJrKksgw0)

Movie S4. Environment-mediated role switching during maze navigation (https://youtu.be/3h1gr8iFp40)

Movie S5. Elastic cargo and interaction across cargo transport collective units (https://youtu.be/Fk51tSPoRWw)

Movie S6. Cooperative transport prefers in-phase; tug-of-war prefers phase lag (https://youtu.be/5Fgyw7ybwaE)

Movie S7. Reversing phase-lag advantage: pull-and-rest, pull-and-brake, or pull-and-hold. (https://youtu.be/Kjr02xKSZYU)

